# The formin FMNL2 plays a role in the response of melanoma cells to substrate stiffness

**DOI:** 10.1101/2024.12.13.628356

**Authors:** Joshua D. Clugston, Sarah Fox, James L. Harden, John W. Copeland

## Abstract

**Background:** Cells are constantly sensing and responding to changes in their local environment to adapt their behaviour and morphology. These external stimuli include chemical and mechanical signals, and much recent work has revealed the complexity of the cellular response to changes in substrate stiffness. We investigated the effects of substrate stiffness on the morphology and motility of A2058 human melanoma cells. FMNL2, a formin protein associated with actin cytoskeleton dynamics, regulates cell morphology and motility but its role in stiffness sensing remains unclear. This study examines how A2058 cells respond to substrates of varying stiffness and evaluates the impact of FMNL2 depletion on these responses.

**Results:** We found that with increasing substrate stiffness the cells transitioned from a rounded cell morphology to progressively more elongated morphologies with a concomitant increase in actin stress fiber alignment. Depletion of FMNL2 expression amplified these morphological changes, with knockdown cells showing consistently greater elongation and more pronounced stress fiber alignment compared to controls. Notably, the orientational order parameter (S) revealed higher alignment of actin filaments along the cell’s long axis in knockdown cells.

Substrate stiffness also affected cell motility, indicated by an apparent optimal stiffness that maximized motility followed by a notable decrease in distance travelled during cell migration on progressively stiffer substrates. This decrease was largely attributable to a decrease in the time the cells spent in motion as the substrate stiffness increased. FMNL2 depletion significantly exacerbated this effect, with knockdown cells traveling shorter net distances and spending less time moving across all substrates.

**Conclusions:** This study demonstrates that substrate stiffness profoundly influences A2058 melanoma cell morphology and motility, with FMNL2 playing a pivotal regulatory role. Our observations suggest that FMNL2 is critical for maintaining motility and morphological adaptability under increased stiffness. Loss of FMNL2 enhanced stress fiber alignment and cell elongation while impairing motility, particularly on stiff substrates, revealing FMNL2 as a mechanosensitive effector. However, further biochemical work should be performed to determine the exact mechanisms by which FMNL2 participates in regulation of melanoma cell response to substrate stiffness.

## I. INTRODUCTION

Directional cell migration is promoted by structural, chemical and physical signals that activate intracellular signaling pathways which govern cytoskeletal dynamics [1]. Indeed, the mechanical properties of the cellular microenvironment have a significant effect on many aspects of cell behaviour, affecting 2D and 3D cell migration as well as cell morphology [2, 3]. Mechanical signals such as substrate stiffness are sensed by actin-based structures that are used to probe the surrounding environment during physiological and pathophysiological migratory events, such as directional migration in response to stiffness gradients (durotaxis [2]). Cell morphology has been shown to be directly linked to substrate stiffness, as cell area often increases on stiffer substrates due to the increased rate of focal adhesion (FA) formation [3–6]. The formation of FAs is often dependent on the generation of tensile forces with the substrate, and the magnitude of this force can change depending on the substrate stiffness [6–9]. On stiffer substrates, cells generate more focal adhesions than on softer substrates, which is often cited as the reason for the increase in cell area as substrate stiffness increases [7, 8]. This response is due, in part, to the cell’s ability to exert higher traction forces, facilitating more significant interactions with the substrate and leading to increased FA formation [7–10]. For these reasons, cell motility is also linked to substrate stiffness, as the increased number of FAs increases the turnover rate, prolonging the cycle of attachment and detachment to the substrate [7, 8]. Mechanosensing is also mediated by the actin stress fibers which are anchored at focal adhesions and directly connect the cytoskeleton to the extracellular environment. Substrate stiffness has also been shown to affect the formation and organization of stress fibers. On rigid substrates, cells will often have well aligned stress fibers and focal adhesions along the major cell axis, while both stress fibers and focal adhesions become radially oriented on softer substrates [11, 12]. In addition to stress fibers, both finger-like filopodia, and flat, sheet-like lamellipodia, have been shown to be part of the cellular apparatus that detects substrate stiffness [2].

During migration, filopodia assist in guiding the cell by probing the environment and establishing connections with the substrate. These protrusions are dependent on formin proteins which directly regulate actin polymerization and the binding and bundling of F-actin [13–17]. Formin like-2 (FMNL2) is a formin protein associated with filopodia assembly in multiple cell lines [17, 18] and is required for filopodia assembly and normal cell motility in melanoma cells and other cancer cell lines [19–23]. Conversely, overexpression of FMNL2 is sufficient to induce filopodia assembly [17, 18, 24]. FMNL2 also contributes to lamellipodia formation and force generation at the leading edge in migrating melanoma cells. Finally, FMNL2 has also been found to be necessary in the formation and turnover of cell-cell contacts, as well as cell-substrate adhesion sites [25, 26]. Thus, FMNL2 activity is involved in the assembly of multiple subcellular structures that are implicated in sensing the physical properties of the extracellular environment.

To shed more light on the role of substrate stiffness in melanoma cell motility, we assessed the behaviour of A2058 human melanoma cells plated on fibronectin functionalized substrates with a range of biologically relevant elastic moduli. We found that A2058 cell morphology changed with increasing stiffness, as the average cell area increased and more elongated cell morphologies with long, well-aligned stress fibers were observed. We also found that these cells exhibit a bimodal mode of cell motility, punctuated by alternating periods of motion and cell arrest. In particular, cell motility peaked at an elastic modulus of 0.5kPa and became progressively less motile on the stiffer substrates, an effect attributable to a reduction in time spent moving on stiffer substrates. We also studied the effect of silencing the protein FMNL2 in A2058 cells. The FMNL2 knockdown cells showed qualitatively similar behaviour, but they were less motile and more elongated than control cells on the same substrate. Depleting FMNL2 expression resulted in the cells spending less time in motion on the stiffer substrates than control cells, which resulted in the average net distance travelled being smaller than the control cells. Our results are consistent with other studies in the literature that show trends for adherent cells to become more elongated and less motile with increasing substrate stiffness. Moreover, the results of this study indicate that the suppression of FMNL2 plays a significant role in modulating cell morphology and motility on substrates of different moduli, implying a potential role for FMNL2 in this process.

## II. RESULTS

### A. Morphological changes across substrates of increasing moduli

Cell proliferation, invasion, and speed have all been shown to change in melanoma cells when they are introduced into environments of different stiffness [25]. To gain insight into this phenomenon, we first characterized the effects of substrate stiffness on the morphology of adherent A2058 human melanoma cells. On extremely stiff substrates (*>*1GPa), such as glass or tissue culture plastic, A2058 cells have a trapezoidal morphology and exhibit extensive stress fiber formation [17]. To test the effects of more physiologically relevant substrate stiffnesses on A2058 morphology we employed substrates with a range of elastic moduli, and to assess a potential role for FMNL2 in this process, FMNL2 expression in A2058 cells was knocked down using siRNA. Control and FMNL2-depleted A2058 cells were plated on fibronectin functionalized silicone gel substrates with moduli of 0.2kPa, 0.5kPa, 2.0kPa, 8.0kPa, and 64.0kPa. On soft substrates (0.2 & 0.5 kPa), control cells adopted a more circular morphology. However, with increasing modulus (2, 8 & 64 kPa), the cells progressively became more elongated (Figure 1A and Table 1). This behavior is reflected quantitatively by the average roundness value, R, which was found to decrease as the substrate modulus increased (Figure 1C). R for the control A2058 cells decreased from 0.763 ± 0.002 on the 0.2kPa substrate to 0.615 ± 0.001 on the 64.0 kPa substrate. A similar trend was observed in the FMNL2 k/d cells (Figure 1B, C), although these cells adopted more elongated conformations in comparison to the control cells for all elastic moduli, with R decreasing from 0.714 ± 0.002 on the 0.2kPa substrate to 0.534 ± 0.001 on the 64.0kPa substrate. Similar results were obtained in FMNL2 k/d cells using a second siRNA duplex (D2), as shown in supplemental Figure S1.

Comparing the R values for the control and knockdown cells on equivalent substrates in Table 1, we can see that their difference, *δR* = *R*_*kd*_ − *R*_*c*_, was consistently negative, indicating that the knockdown cells adopted more elongated morphologies then their control counterparts. The cell area (A), perimeter (P), and Feret diameter (FD), defined as the longest distance between two points along an object’s boundary, were also calculated for each cell and averaged over the cell population to further characterize the effect of substrate stiffness on cell morphology. All values were found to monotonically increase with increasing substrate stiffness, as shown in Table 1. For insight, we examined the net average change of these parameters (Δ*A*, Δ*P*, Δ*FD*, and Δ*R*) relative to their values on the softest substrate (0.2kPa), listed in Table 2.

**TABLE I:**
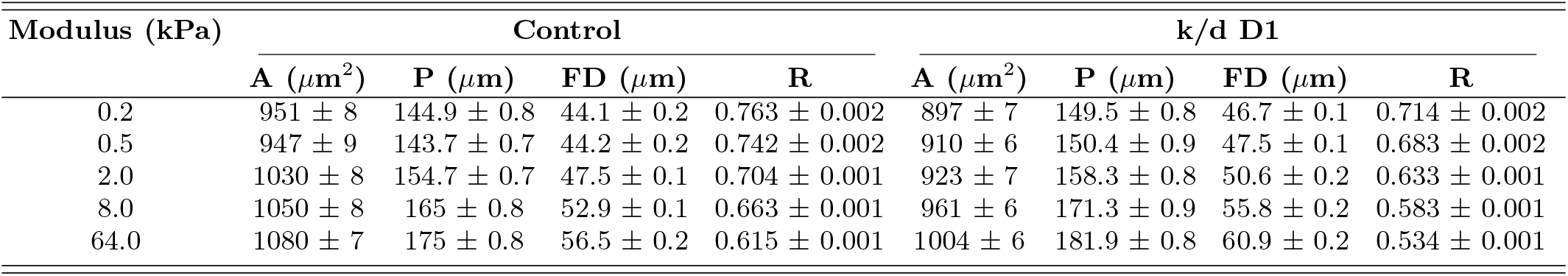
Morphological changes in A2058 cells on increasing moduli substrates for control and FMNL2 depleted cells. The average area (A), perimeter (P), Feret diameter (FD), and Roundness (R) are shown for the control and knockdown (k/d) cells on each substrate. The error on each value is the standard error of the mean.

**TABLE II:** Net morphological changes between A2058 cells on all substrates relative to the 0.2kPa substrate, \**p <* 0.0001.

| Modulus (kPa) | Control |  |  |  | k/d D1 |  |  |  |
| --- | --- | --- | --- | --- | --- | --- | --- | --- |
| | $\Delta A$ ($\mu\text{m}^2$) | $\Delta P$ ($\mu\text{m}$) | $\Delta FD$ ($\mu\text{m}$) | $\Delta R$ | $\Delta A$ ($\mu\text{m}^2$) | $\Delta P$ ($\mu\text{m}$) | $\Delta FD$ ($\mu\text{m}$) | $\Delta R$ |
| 0.5 | $-4 \pm 10$ | $-1.2 \pm 1.1$ | $0.1 \pm 0.2$ | $-0.021 \pm 0.003^*$ | $13 \pm 9$ | $0.9 \pm 1.2$ | $0.8 \pm 0.1^*$ | $-0.031 \pm 0.003^*$ |
| 2.0 | $79 \pm 11^*$ | $9.8 \pm 1.1^*$ | $3.4 \pm 0.2^*$ | $-0.059 \pm 0.002^*$ | $26 \pm 10^*$ | $8.8 \pm 1.1^*$ | $3.9 \pm 0.2^*$ | $-0.081 \pm 0.002^*$ |
| 8.0 | $99 \pm 11^*$ | $20.1 \pm 1.1^*$ | $8.8 \pm 0.2^*$ | $-0.100 \pm 0.002^*$ | $64 \pm 9^*$ | $21.8 \pm 1.2^*$ | $9.1 \pm 0.2^*$ | $-0.131 \pm 0.002^*$ |
| 64.0 | $129 \pm 11^*$ | $30.1 \pm 1.1^*$ | $12.4 \pm 0.3^*$ | $-0.148 \pm 0.002^*$ | $107 \pm 9^*$ | $32.4 \pm 1.1^*$ | $14.2 \pm 0.2^*$ | $-0.180 \pm 0.002^*$ |

The average changes in area ( $\Delta A$ ), perimeter ( $\Delta P$ ), Feret diameter ( $\Delta FD$ ), and Roundness ( $\Delta R$ ) are shown for control and knockdown (k/d) cells on each substrate. Asterisks (\*) indicate statistically significant differences ( $p < 0.0001$ ).

**FIG. 1:**
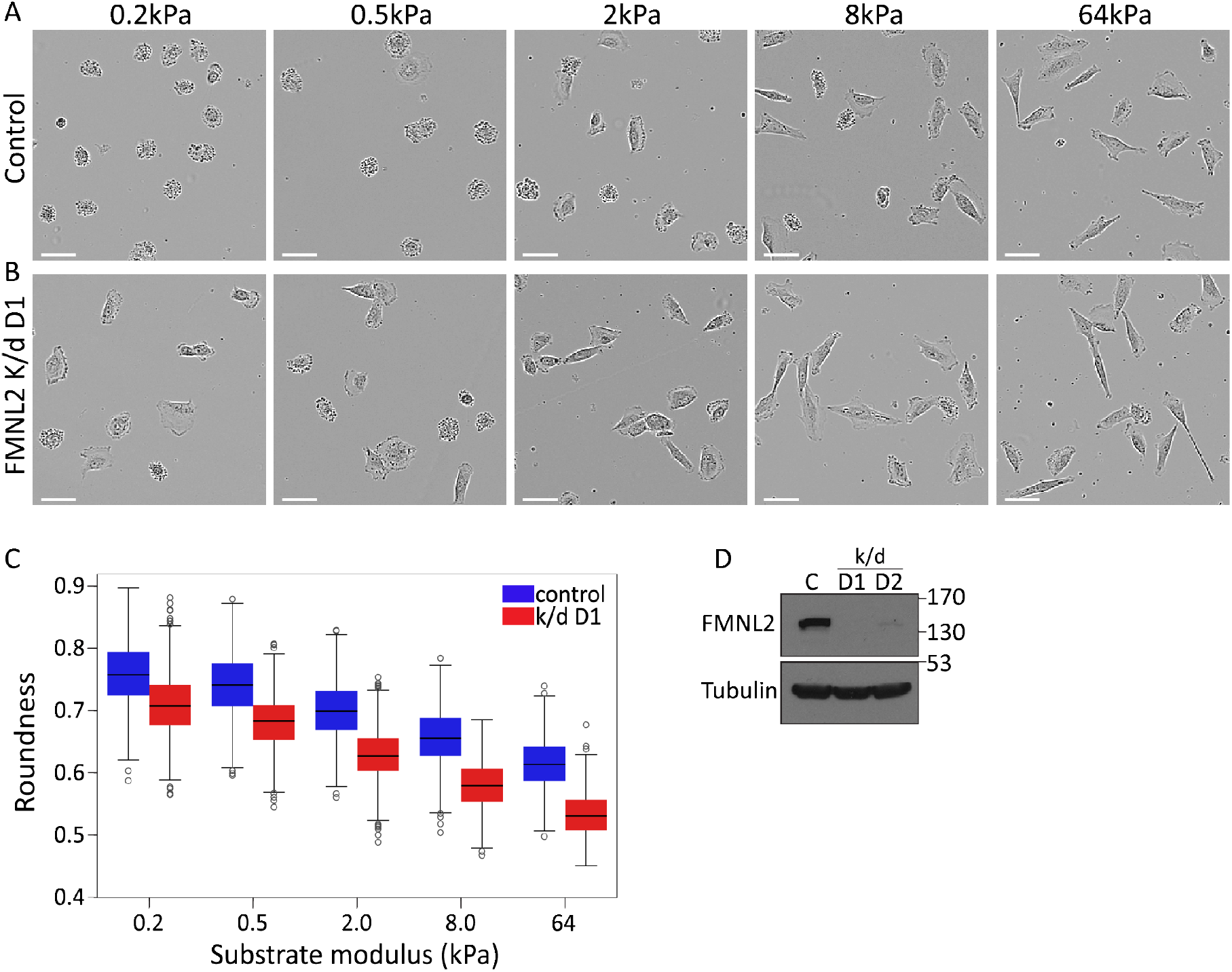
Figure 1. A2058 Morphology Changes Across Substrates with Increasing Modulus. (A) Representative phase contrast images of control A2058 cells plated on substrates with the indicated elastic moduli. (B) Representative phase contrast images of FMNL2-depleted A2058 cells plated on substrates with the indicated moduli. Scale bar=25 mm. (C) The average roundness value R for control and FMNL2 knockdown cells across all substrates (see Table 1 for all cell morphology parameter values, *p*<*0.0001). ° indicates values distributed outside of 2 standard deviations. N=3 (D) Anti-FMNL2 immunoblot confirming the extent of FMNL2 knockdown by transient transfection of siRNA. Duplex D1 was consistently more effective in FMNL2 depletion than duplex D2. Tubulin was used as a loading control.

For instance, the net average area increase, Δ*A*, between the 0.2kPa substrate and the 64.0kPa substrate was Δ*A* = 129 *±* 11*µm*^2^ for the native A2058 cells and Δ*A* = 107 *±* 9*µm*^2^ for the FMNL2 k/d cells. Like-wise, the net average cell perimeter and Feret diameter increases between the 0.2kPa and 64.0kPa substrates, with Δ*P* = 30.1 *±* 1.1*µm* and Δ*FD* = 14 *±* 0.3*µm* for the control cells and with Δ*P* = 32.4 *±* 1.1*µm* and Δ*FD* = 14.2 *±* 0.2*µm* for the FMNL2 k/d cells (Table 2). This increase in area and perimeter indicates that the control cells were more spread as the substrate stiffness increased, while the increase in Feret diameter shows that these cells also became more elongated with increasing stiffness. Notably, when compared to the control A2058 cells, FMNL2 k/d cells showed smaller values of average cell area (A), larger values of average cell perimeter (P), and larger values for Feret diameter (FD) on equivalent stiffness substrates. Table 3 reports the differences (*δA, δP, δFD*, and *δR*) between values of A, P, FD, and R for FMNL2 k/d cells relative to the control cells. The observation of negative *δA* and positive *δP* and *δFD* for different modulus substrates further corroborates the negative relative average roundness *δR* results, as these changes indicate that the knockdown cells became more elongated than the control cells on the same substrate. Similar results were obtained in FMNL2 k/d cells using a second siRNA duplex (D2), as shown in supplemental Tables S1-S3.

**TABLE III:**
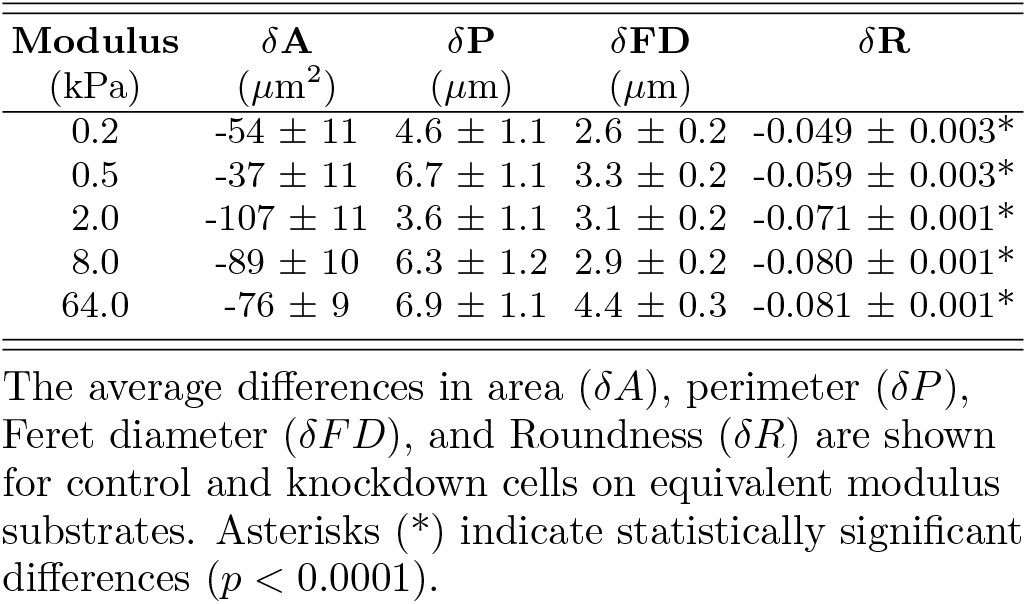
Net morphological changes between A2058 control and FMNL2 depleted cells on equivalent modulus substrates, \**p <* 0.0001.

| Modulus (kPa) | $\delta A$ ($\mu\text{m}^2$) | $\delta P$ ($\mu\text{m}$) | $\delta FD$ ($\mu\text{m}$) | $\delta R$ |
| --- | --- | --- | --- | --- |
| 0.2 | $-54 \pm 11$ | $4.6 \pm 1.1$ | $2.6 \pm 0.2$ | $-0.049 \pm 0.003^*$ |
| 0.5 | $-37 \pm 11$ | $6.7 \pm 1.1$ | $3.3 \pm 0.2$ | $-0.059 \pm 0.003^*$ |
| 2.0 | $-107 \pm 11$ | $3.6 \pm 1.1$ | $3.1 \pm 0.2$ | $-0.071 \pm 0.001^*$ |
| 8.0 | $-89 \pm 10$ | $6.3 \pm 1.2$ | $2.9 \pm 0.2$ | $-0.080 \pm 0.001^*$ |
| 64.0 | $-76 \pm 9$ | $6.9 \pm 1.1$ | $4.4 \pm 0.3$ | $-0.081 \pm 0.001^*$ |

The average differences in area ( $\delta A$ ), perimeter ( $\delta P$ ), Feret diameter ( $\delta FD$ ), and Roundness ( $\delta R$ ) are shown for control and knockdown cells on equivalent modulus substrates. Asterisks (\*) indicate statistically significant differences ( $p < 0.0001$ ).

### B. Substrate stiffness impacts organization of actin filaments

To further assess the changes in cell morphology, control and FMNL2 k/d cells plated on substrates of increasing stiffness were fixed and stained with phalloidin to image filamentous actin. The orientation and structure of the actin fibers were notably different between the soft and stiff substrates in both control and knockdown cells (Figure 2A-C). To quantify this effect, we defined a 2D orientational order parameter (S), which quantifies the alignment of the actin fibers within the cell with respect to a defined axis - the direction of the cell elongation as given by the Feret diameter. This value can vary between -1 and 1, with values close to 0 indicating random orientation, values close to 1 showing preferential alignment of the stress fibers within the cell along the Feret diameter, and values close to -1 indicating alignment perpendicular to the Feret diameter. As the substrate modulus increased, we found that S for both the control and knockdown cells increased with substrate modulus (Figure 2A), and the angular orientation of the fibers became more tightly distributed around 0^*°*^, i.e. parallel to the direction of the Feret diameter (Figure 2E). Furthermore, there was a noticeable enhancement of S in the knock-down cells relative to the control cells. Moreover, the angular distribution of the knockdown cell fibers was more closely distributed around 0^*°*^ for the knockdown cells. Similar results were obtained with knockdown of FMNL2 using a second siRNA duplex, D2 (Supplemental Figure S2). The values of S and the width of the distributions of fiber orientation both indicate that the actin fibers in the FMNL2 k/d cells were consistently more aligned with the long axis of the cell in comparison to the control cells.

**FIG. 2:**
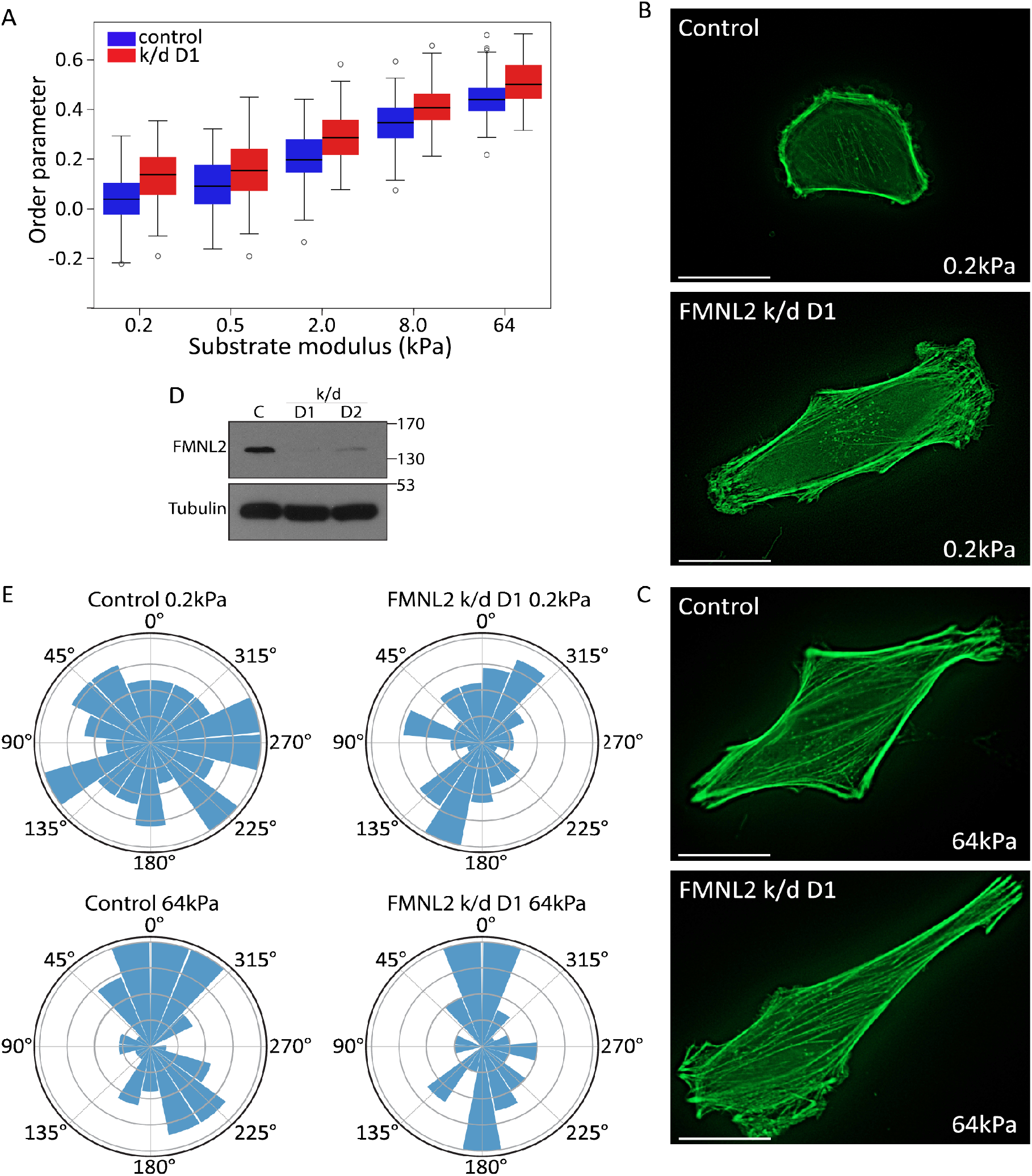
Figure 2. Substrate modulus affects actin stress fiber orientation in A2058 cells. (A) 2D orientational order parameter S quantifying average orientation of F-actin with respect to the cell Feret diameter. Increasing substrate stiffness increases the 2D order parameter. ° indicates values outside of 2 standard deviations. N=3 (B, C) Representative images of the cells analyzed in (A) for control and FMNL2 depleted cells plated on the indicated substrates. Cells were fixed and stained with phalloidin to visualize actin filaments. Scale bar=20 mm (D) Anti-FMNL2 immunoblot confirming the extent of FMNL2 knockdown by transient transfection of siRNA. Tubulin was used as a loading control. (E) Windrose plots showing the orientation of the actin fibers with respect to the cells’ Feret diameter for control and FMNL2 depleted cells plated on 0.2kPa and 64kPa substrates. Increases in substrate modulus and FMNL2 depletion both yield a tighter distribution around 0, indicating increased filament alignment.

### C. Changes in cell motility with increasing substrate modulus

To gain further insight into the effects of substrate stiffness on cell behaviour, A2058 cells were tracked for 24 hours to compare cell motility across the different substrates. These cells exhibited a bimodal pattern of cell motility characterized by alternating periods of motion and arrest. Overall, for both control and FMNL2 k/d A2058 cells, the average net distance travelled, 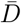, decreased with increasing substrate modulus (Figure 3). Across all five substrates tested, the knockdown cells consistently travelled an average net distance 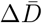between 39 *±*3*µm* and 55 *±*4*µm* less than the control cells (Table 4). This is clearly observed in the shift in distribution of distance D travelled for the knockdown cells towards a smaller average value across all substrates (Figure 3A-E).

**TABLE IV:** The average net distance traveled for control and FMNL2 depleted cells.

| Modulus<br>(kPa) | Control<br>$\bar{D}$ ($\mu\text{m}$) | k/d D1<br>$\bar{D}$ ($\mu\text{m}$) | Difference<br>$\Delta\bar{D}$ ($\mu\text{m}$) |
| --- | --- | --- | --- |
| 0.2 | $329 \pm 4$ | $281 \pm 4$ | $-48 \pm 6^*$ |
| 0.5 | $340 \pm 4$ | $293 \pm 4$ | $-47 \pm 6^*$ |
| 2.0 | $303 \pm 4$ | $264 \pm 4$ | $-39 \pm 6^*$ |
| 8.0 | $282 \pm 3$ | $238 \pm 4$ | $-49 \pm 5^*$ |
| 64.0 | $242 \pm 3$ | $187 \pm 3$ | $-55 \pm 4^*$ |

Table values for the average net distance traveled ( $\bar{D}$ ) for control and FMNL2 depleted cells and the relative values ( $\Delta\bar{D}$ ) on equivalent substrates. $*p < 0.0001$ , $N=3$ .

**FIG. 3:**
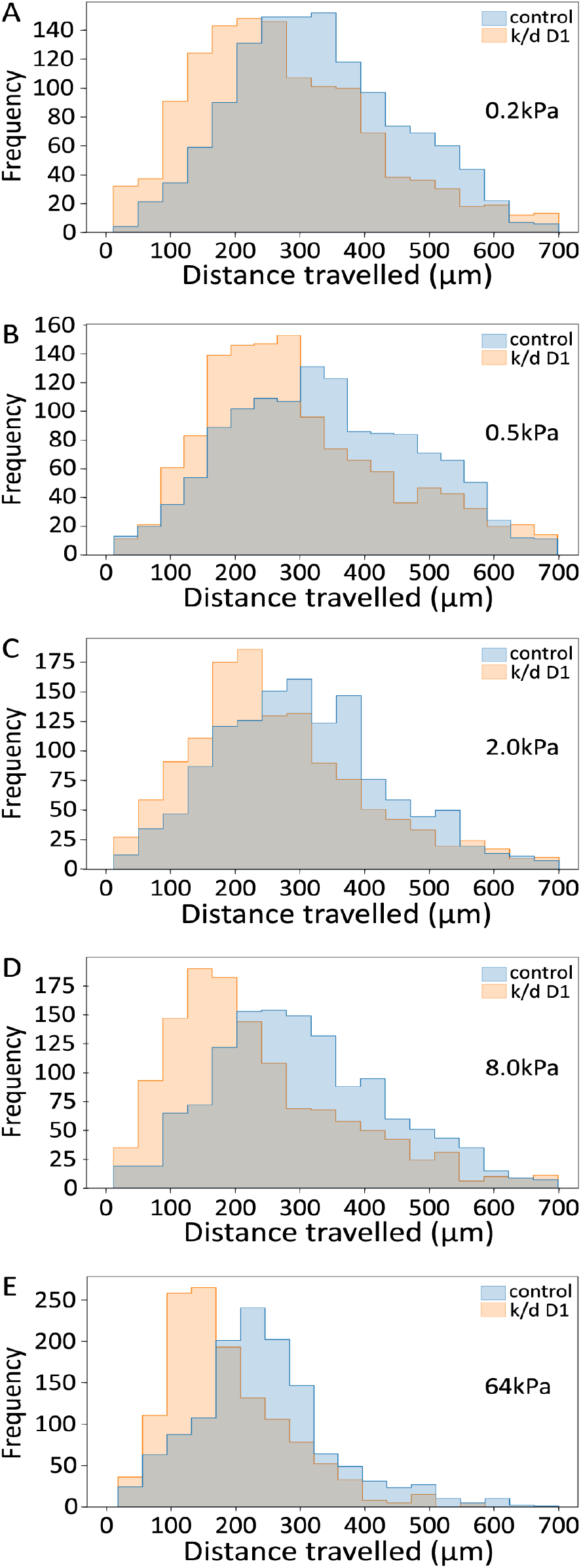
Figure 3. Cell motility is decreased with increased modulus. (A-E) Histograms of the net distance D travelled across all substrates for control (blue) and FMNL2 knockdown (orange) A2058 cells, with the overlap region shown in gray. Knockdown cells show a smaller average distance travelled than control cells as indicated by the shift in the histogram. Values are derived from cells tracked by live-cell imaging from the experiments shown in figure 1.

To characterize the punctuated cell motility observed for A2058 cells, we next examined the ensemble-averaged time spent moving by the cells, *t*_*m*_, during a motility experiment and its dependence on substrate stiffness. Moreover, the ensemble-averaged cell speed *s*_*m*_ was also calculated using *t*_*m*_ and 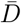, and correlated with substrate modulus. This comparison can determine whether cell motility is intrinsically slower on stiffer substrates, or rather if cells spend progressively less time moving on substrates of increasing stiffness. It also more clearly assesses the underlying differences in motility between the control and knockdown cells. The average time in motion *t*_*m*_ for both control and FMNL2 k/d cells first increased slightly from the 0.2kPa to the 0.5kPa modulus substrates, before monotonically diminishing with increasing substrate stiffness above 0.5kPa (Figure 4A and Table 5). Moreover, the average cell moving speed *s*_*m*_ followed these same trends with increasing stiffness (Table 6) for both groups, but in a substantially subdued manner. Interestingly, the FMNL2 k/d cells spent significantly less time in motion than the control cells on the 2.0kPa, 8.0kPa, and 64.0kPa substrates, while the average time in motion was found to be statistically equivalent between control and knockdown cells on the 0.2kPa and 0.5kPa substrates, as shown by Δ*t*_*m*_ in Table 5. Comparing the average moving speed *s*_*m*_ between the two cell types on equivalent substrates, the knock-down cells moved more slowly than the control A2058 cells, but the differences, as shown by Δ*s*_*m*_ in Table 6, are less pronounced than those seen in the time spent moving. Depletion of FMNL2 expression with the second siRNA duplex produced similar results (Supplemental Figure S4). Overall, the decrease seen in the average distance travelled by both cell types with increasing substrate modulus is largely attributable to their decreased time spent in motion, with a smaller contribution from their decreased average speed, with FMNL2 k/d cells moving systematically less than control A2058 cells on equivalent substrates.

**TABLE V:** The average time spent in motion for control and knockdown cells.

| Modulus<br>(kPa) | Control | | Difference<br>$\Delta t_m$ (min) |
| --- | --- | --- | --- |
| | $t_m$ (min) | $t_m$ (min) | |
| 0.2 | $561 \pm 2$ | $553 \pm 2$ | $-8 \pm 3^*$ |
| 0.5 | $574 \pm 2$ | $561 \pm 2$ | $-13 \pm 3^{****}$ |
| 2.0 | $543 \pm 2$ | $504 \pm 2$ | $-39 \pm 3^{****}$ |
| 8.0 | $501 \pm 2$ | $453 \pm 2$ | $-48 \pm 3^{****}$ |
| 64.0 | $483 \pm 2$ | $415 \pm 2$ | $-68 \pm 3^{****}$ |

Values for the average time spent moving ( $t_m$ ) for control and FMNL2 knockdown cells and the difference in the average time spent moving ( $\Delta t_m$ ) between control and knockdown cells on the same substrate.
\* $p < 0.05$ , \*\* $p < 0.01$ , \*\*\* $p < 0.001$ , \*\*\*\* $p < 0.0001$ .

**TABLE VI:** The average moving cell speed for control and knockdown A2058 cells.

| Modulus<br>(kPa) | Control | | Difference<br>$\Delta s_m$ ($\mu\text{m}/\text{h}$) |
| --- | --- | --- | --- |
| | $s_m$ ($\mu\text{m}/\text{h}$) | $s_m$ ($\mu\text{m}/\text{h}$) | |
| 0.2 | $35.5 \pm 0.2$ | $34.9 \pm 0.3$ | $-0.6 \pm 0.4^{***}$ |
| 0.5 | $36.1 \pm 0.8$ | $35.4 \pm 0.3$ | $-0.7 \pm 0.4^{****}$ |
| 2.0 | $32.7 \pm 0.3$ | $30.9 \pm 0.8$ | $-1.8 \pm 0.4^{****}$ |
| 8.0 | $33.6 \pm 0.3$ | $30.8 \pm 0.3$ | $-2.8 \pm 0.4^{****}$ |
| 64.0 | $29.6 \pm 0.2$ | $28.4 \pm 0.3$ | $-1.2 \pm 0.4^{****}$ |

Average cell speed ( $s_m$ ) for control and knockdown cells and the difference in the average cell speed ( $\Delta s_m$ ) between control and knockdown cells on the same substrate. For both control and FMNL2 knockdown cells there is a decrease in the average moving speed as the modulus increases. \*\* $p < 0.01$ , \*\*\* $p < 0.001$ , \*\*\*\* $p < 0.0001$ .

**FIG. 4:**
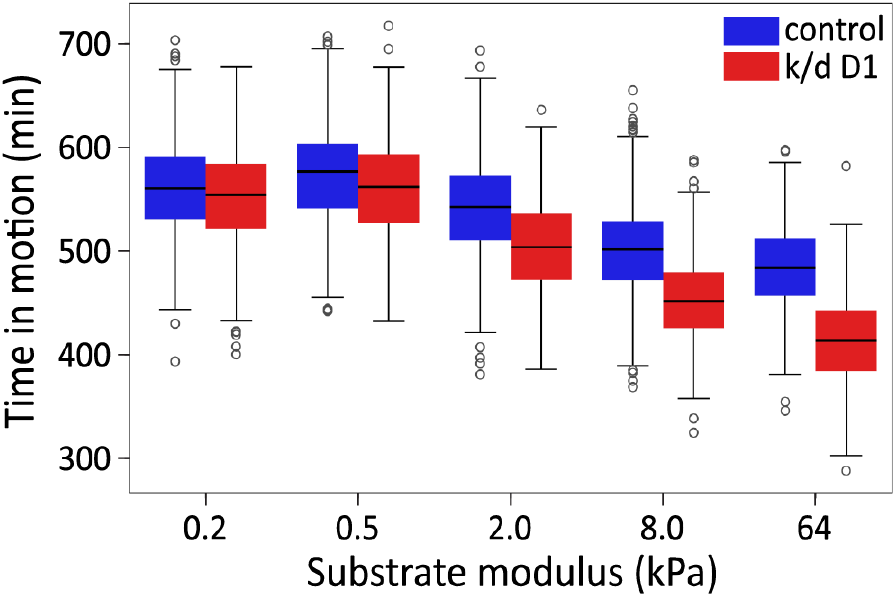
Figure 4 — Time spent in motion for control and FMNL2 knockdown cells. A box plot showing the time spent moving for control (blue) and FMNL2 knockdown (red) cells with increasing substrate modulus. Increasing stiffness resulted in decreased time in motion. FMNL2 depleted cells spent less time in motion that control cells on each substrate, N=3.

## III. DISCUSSION

In this study, we tested the effects of substrate stiffness on the morphology and motility of A2058 human melanoma cells. On the softest substrates, the cells consistently adopted circular morphologies and as the substrate stiffness increased, they became progressively more elongated. This effect was quantified by the cell roundness parameter R which confirmed the impact of substrate stiffness on cell shape. The increase in elongation (decrease in R) was matched by an increase in the average area A, perimeter P and Feret diameter FD of each cell, consistent with the cells becoming more elongated and more spread as the substrate became stiffer. Similarly, the increase in elongation on stiffer substrates was accompanied by an increase in stress fiber formation and stress fiber alignment as quantified by the rising fibre orientational order parameter S. Knockdown of FMNL2 expression diminished the increase in cell area with increasing stiffness relative to the control cells and was associated with a relative increase in cell elongation and actin filament alignment with increasing stiffness. Cell motility is driven by forces generated by the actin network pushing against the lamellipodial membrane at the leading edge and also by traction forces generated by integrin adhesion complexes (IAC). Molecular clutch theory suggests that cells move toward regions where the highest force is exerted. In positive durotaxis, higher force is generated on stiffer substrates and is dependent on the focal adhesion protein Talin acting as part of the IAC mechanosensory apparatus [27]. In other cases, cells migrate toward an optimal stiffness determined by the substrate modulus and the cytoskeletal regulatory machinery [2]. In such cases, negative durotaxis may occur, in which cells migrate towards softer substrates where an optimal stiffness generates maximum force. Indeed, we found that A2058 cells exhibited two distinct motility regimes: an initial regime where cell motility first increased modestly with increasing elastic modulus below 0.5kPa, followed by a second regime where cells became significantly less motile with further increase of elastic modulus. We also note a correlation between the substrate induced changes in cell roundness and motility in A2058 cells in both regimes. On the softer substrates 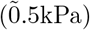, cells exhibited a rounder morphology and enhanced motility, while on stiffer substrates cells became progressively elongated and exhibited diminishing motility with increasing modulus, suggesting the existence of an optimal stiffness for the migration of these cells near an elastic modulus of 0.5kPa. The behaviour observed for A2058 cells on the softer substrates is consistent with previous reports of cells that undergo positive durotaxis [11, 12]. In contrast, on stiffer substrates the changes in A2058 cell motility with increasing modulus are more consistent with cells that exhibit negative durotaxis [2, 27]. Further analysis revealed that with increasing substrate stiffness above 0.5kPa, the decrease in cell motility was largely attributable to a substantial decrease in the average time in motion of the cells, with the concomitant decrease in average speed while in motion playing a relatively minor role. As with the effects on cell morphology, FMNL2 depletion also inhibited cell motility across all substrates with a greater effect on stiffer substrates (2.0, 8.0 and 64 kPa. As with WT A2058 cells, the decrease in the motility of the FMNL2 knockdown cells was largely due to the decrease in the average time in motion of the cells with increasing substrate modulus, possibly providing clues to the biological origins of this motility inhibition.

FMNL2 plays an important role in facilitating melanoma cell motility [18, 24, 28] and we found the FMNL2 knockdown inhibited cell migration on all the substrates tested. This effect was significantly more pronounced on the stiffer substrates suggesting that the function of FMNL2 in cell motility is governed by substrate stiffness. FMNL2 knockdown also had a significant effect on the stiffness induced changes in morphology. The knockdown cells were more elongated and had appreciably more stress fibers aligned with the long axis of the cell. Enhanced stress fiber alignment was somewhat unexpected given the association of formins with the assembly of these structures [29]. Similar results were obtained, however, following the depletion of the related protein FMNL1 in other cell-types [30] suggesting this might be a reflection of the function of this formin subfamily in the regulation of actin dynamics. Indeed, FMNL2 activity is more closely associated with filopodia formation [13–18, 31] or with force generation at lamellipodia [24, 32]. FMNL2 is also found in the melanoma cell adhesome [33]. Each of these structures has been previously suggested to acts as part of the mechanosensory apparatus [2, 34]. This has interesting implications for the role of FMNL2 in the cellular response to substrate stiffness. Is FMNL2 part of the mechanosensing apparatus acting either at filopodia, lamellipodia or the IAC, or is it a downstream effector that is regulated by mechanosignaling? Our results do not test this directly but are more consistent with an effector role for FMNL2 given its more modest effects on cell behaviour on the softest substrates.

## IV. CONCLUSION

In summary, we found that increasing substrate stiffness above a characteristic value (near 0.5kPa) had a marked effect on the morphology and motility of A2058 melanoma cells. These cells were observed to become move elongated, travel shorter distances, and spend less time in motion as the substrate modulus increased, behaviour consistent with cells that undergo negative durotaxis. In contrast, cell motility was somewhat enhanced with increasing stiffness below 0.5kPa, consistent with an optimal substrate modulus near 0.5kPa and positive durotaxis below this value. Interestingly, cell migration was inhibited in FMNL2 knockdown cells in comparison to control cells on the same substrates, indicating systematic quantitative differences between WT and knockdown cells. These findings highlight the significant impact of substrate stiffness on A2058 cell behavior and suggest that FMNL2 plays an important role in the pathways governing the response of A2058 cells to their extracellular environment.

## V. METHODS

### A. Cell Culture

A2058 (CRL-11147) melanoma cells obtained from the American Type Culture Collection were cultured in Dulbecco’s modified Eagle’s medium (Wisent; 319-007 CL) supplemented 10% v/v with fetal bovine serum (Wisent; 090-150-FBS) in 5% CO2 according to the supplied guidelines. Mycoplasma contamination was tested biweekly. Advanced Biomatrix CytoSoft^®^ 6-well Plates– discovery kit #5190: Eppendorf 6-well cell culture plate (Eppendorf 0030.720.113) with a 0.5mm layer of activated biocompatible silicone of defined elastic modulus were used for all live-cell imaging. Advanced Biomatrix CytoSoft^®^ Imaging 24-well Plate 0.2kPa (#5183) and 64kPa (#5189): Eppendorf 24 well cell imaging plate (0030.741.021) #1.5 glass bottom with a 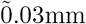 layer of activated biocompatible silicone of defined elastic modulus were used for fixed cell imaging. Silicone surfaces were coated with a Fibronectin solution (bovine plasma, Sigma; F1141) at a final concentration of 10µg/ml in DPBS (Wisent; 311-425 CL) for 1 hour at room temperature. The fibronectin solution was removed, and the plates maintained in DPBS until the cells were added. siRNA-mediated knockdown was performed as previously described [17] using Dharmafect1 (Horizon Discovery Ltd; T-2001-03) and the following siRNA duplexes: FMNL2 siRNA Duplex1 (IDT; hs.Ri.FMNL2.13.1); FMNL2 siRNA duplex2 (IDT; hs.Ri.FMNL2.13.2).

### B. Microscopy

Live cell imaging was performed on an Incucyte^®^ S3 Live-Cell Analysis System (Sartorius). Full technical specifications can be found here: https://www.sartorius.com/download/930502/incucyte-s3-technical-specification-sheet-8000-0527-c00-en-s-1-data.pdf. High-definition phase-contrast images were acquired using the 10X/NA 0.3 objective with an image resolution of 1.2µm/pixel. 72h after transfection with control or FMNL2 siRNA duplex, A2058 cells were seeded at a density of 60, 000 cells per well in CytoSoft^®^ 6-well Plates (Advanced Biomatrix – discovery kit #5190) with the indicated moduli ranging from 0.2kPa to 64kPa. Remaining cells were transferred to fresh 3.5cm dishes and incubated at 37°C, 5% CO2. Cells were imaged using the Incucyte^®^ S3 Live-Cell Analysis System at 37°C in a 5% CO2 incubator. Phase images were collected every 20 minutes for 24h, the first time-point at 2.5h post seeding. Cells were then fixed for 10 minutes with freshly prepared 4% paraformaldehyde in PHEM (PIPES, Hepes, EGTA, MgCl2) for additional analysis [35]. The parallel cell samples were harvested and boiled in 1X Laemmli buffer to assess knockdown efficiency by immunoblotting.

High resolution fixed cell images were acquired using a Zeiss Axio Observer 7 inverted microscope with linear encoded stage and HXP 120V light source with built in power supply, shutter, lamp module and infra-red filter. Zeiss filter set 37 Ex. BP 450/50 FT:480 Em. BP 510/50, 63X 1.4NA oil immersion Plan-apochromat objective, detection with a Hamamatsu ORCA-Flash LT 16bit camera. Z-stacks captured with Zeiss Zen 3.0 software. 55h after transfection with control or FMNL2 siRNA duplex, A2058 cells were seeded at a density of 2, 250 cells per well in CytoSoft^®^ Imaging 24-well plates 0.2kPa and 64kPa moduli and incubated at 37°C, 5% CO2. Remaining cells were transferred to fresh 3.5cm dishes and incubated at 37°C, 5% CO2. 72h post transfection, cells were fixed for 10 minutes with freshly prepared 4% paraformaldehyde in PHEM (PIPES, Hepes, EGTA, MgCl2) [35]. The parallel cell samples were harvested in 1X Laemmli buffer and assessed for knockdown efficiency by immunoblotting.

### C. Immunofluorescence

Immunofluorescence was performed as in [17]. Briefly, cells fixed in 4% formaldehyde/PHEM buffer were permeabilized and blocked for 20 min in 0.3% Triton X-100, 5% donkey serum in 1× PBS, washed in 1× PBS, and incubated with Alexa Fluor 488 Phalloidin (Molecular Probes^®^; A12379) diluted 1:200 in 0.03% Triton X-100 and 5% donkey serum in 1× PBS for 1 h at room temperature. Washed and stored in 1X PBS.

### D. Immunoblotting

siRNA knockdown efficiency was assessed by immunoblotting. Cells were washed with 1X PBS and harvested in 1x Laemmli buffer. Lysates were subjected to SDS-PAGE and immunoblotted with the indicated antibodies. Chemiluminescence was used for detection using the Immobilon^®^ Crescendo western HRP substrate reagent (Millipore Sigma). FMNL2 was detected using chicken anti-FMNL2 [18] and Peroxidase AffiniPure^™^ Donkey Anti-Chicken IgY (IgG) (H+L) (Jackson ImmunoResearch Laboratories); tubulin with mouse anti-*α*-tubulin (Sigma; T5168) and Peroxidase AffiniPure^™^ Donkey Anti-Mouse IgG (H+L) (Jackson ImmunoResearch Laboratories).

### E. Statistical & error analysis

All experimental measurements were recorded, and the mean and standard deviation (SD) of these values were calculated. The uncertainty associated with each mean value was calculated using the standard error of the mean (SEM). For calculating the uncertainty on measurements using the calculated mean values, propagation of error was used to determine their associated error. To determine statistical significance between two measured values, Analysis of Variance (ANOVA) testing was performed to discern any statistically significant differences between the groups. Tukey’s post hoc analysis was then performed to identify significant results between the mean values of different groups. All statistical analyses were conducted with a pre-established alpha level of 0.05, denoting the threshold for statistical significance.

### F. Morphology & motility analysis

Using the Python package OpenCV, the tiff stacks were first thresholded, a process where the pixels are binarized based on their intensity, with pixels having a value less/greater than the defined threshold assigned to white (0)/black (1). The tiff stacks are subsequently filtered using a Gaussian blur to reduce noise in the image. To quantify the morphology of cells, the cells were first located using the OpenCV function “findContours”, which identifies the boundary of objects in binarized images by looking for sharp increases or decreases in adjacent pixel values. Following this, the function “fitEllipse” was then used to fit an ellipse to the contours identified in the previous step. Using the contours, a mask is then created by converting all the pixels populating the inside of the contour into a binary image. The major and minor axis of the ellipse were then extracted and exported to a csv file, as well as the area, perimeter, and spatial coordinates of the boundary of the mask. With these values, we can calculate the roundness of each cell (*r*_*i*_) as the ratio of the minor to major axis length of the fitted ellipse. We can then average over all N cells on the substrate to obtain the group average roundness,

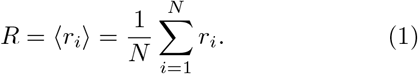

To calculate the Feret diameter, a custom Python script calculates the distance between all points around the boundary of the mask, and then extracts the maximum value.

To track the cells, they were located using OpenCV’s contour finding function, and then tracked for the entire tiff stack with the Discriminative Correlation Filter with Channel and Spatial Reliability (CSRT) tracker in OpenCV. This object tracking algorithm works by applying discriminative correlation filters to different feature channels of the image (color, texture, etc) to determine their reliability. Each channel is weighted independently of the others based on its assigned reliability, which the CSRT tracker assesses by measuring each channel’s consistency in response over time, focusing on signal-to-noise ratio (SNR) to emphasize stable features and suppress noisy ones. The tracker also learns to discriminate between the object and its background to enhance accuracy when the background may contain distracting elements. At each time point, the position of each cell along each the x and y axis were then recorded, and exported as a csv file and analyzed to determine all motility measurements once tracking was complete. The net distance travelled by each cell was then calculated by summing the distance travelled between each time step Δ*t* from an initial time *t*_0_ up to the total time T:

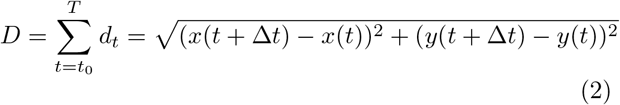

The speed of each cell between each time step was also calculated to determine if the cell was moving. If the speed of the cell was less than 5*µ*m/h, than it was classified as not moving, as speeds less than this were often due to morphological changes in the cell that changed its center of mass, and not true movement. The time spent stationary and the time spent moving were then calculated for each cell using this restriction. These were averaged over the cell population to determine an ensemble-averaged moving time *t*_*m*_. Likewise, the ensemble-averaged speed *s*_*m*_ was determined by averaging the mean cell speed during a cell trajectory over the cell population.

### G. Actin fiber analysis

For actin fiber alignment analysis, images were first preprocessed in ImageJ by enhancing the contrast of the image by 0.35%. Subsequently, the stack was then exported as a maximum projection intensity image. A binary mask was also created and exported by thresholding the image to separate the cell from the background. The analysis was then performed with a previously used method known as Alignment by Fourier Transform [36] with some modifications to their python scripts.

This method segments the windows of a defined size, performs a fast Fourier transform on each window in the image, and then outputs a vector field representing the alignment of F-actin fibers in a cell. Each vector has its orientation angle *θ* with respect to a central reference vector oriented along the major axis of the cell calculated, which is then used to find the local F-actin orientational order parameter of each cell [37]

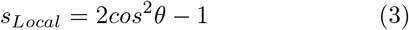

This value describes the orientation of the F-actin with respect to the major axis of the cell. All calculated order parameters are then averaged over one cell to obtain the orientational order parameter of the entire cell

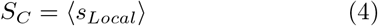

We can then average over all the cells on a given substrate to obtain the global orientational order parameter

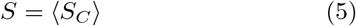

## VI. DECLARATIONS

### Authors’ contributions

JC, JLH and JWC designed the study. JC and SF performed the experiments. JC analysed the data and was the major contributor in writing the manuscript. JLH and JWC supervised the research project. All authors read and approved the final manuscript.

## Acknowledgments

The authors acknowledge the outstanding support of the Cell Biology and Image Acquisition Core (*RRID* : *SCR*_0_21845) at the University of Ottawa, Faculty of Medicine.

## Availability of data and materials

The datasets used and/or analysed during the current study are available from the corresponding author on reasonable request.

## Consent for publication

Not applicable.

## Ethics approval and consent to participate

Not applicable.

## VII. SUPPLEMENTARY FIGURES & TABLES

**FIG. S1:**
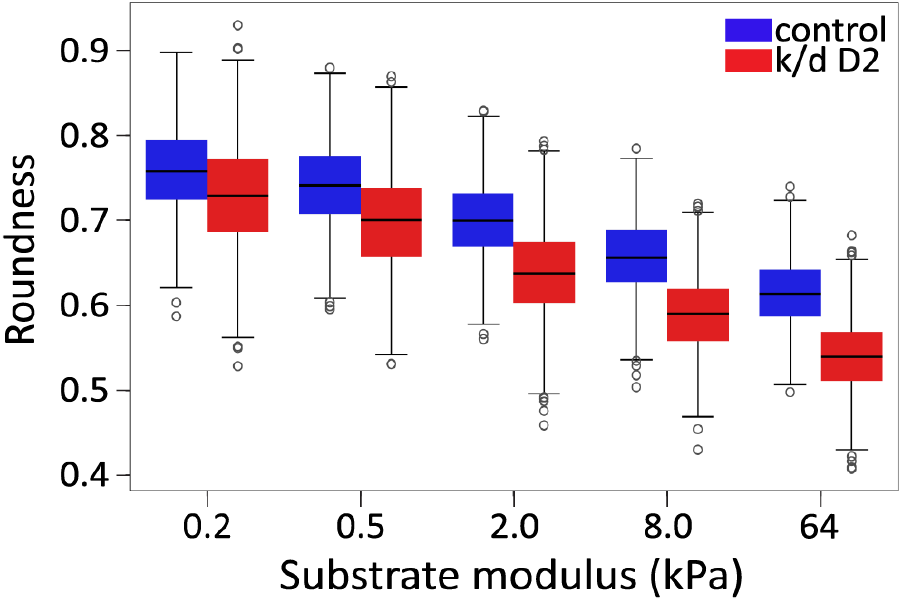
A2058 Morphology Changes Across Substrates with Increasing Modulus. As in Figure 1, the average roundness value *R* for control and FMNL2 knockdown cells across all substrates was calculated based on images captured during live-cell imaging (see additional Table S1 for all siRNA duplex D2 cell morphology parameter values). ° indicates values distributed outside of 2 standard deviations. N=3.

**FIG. S2:**
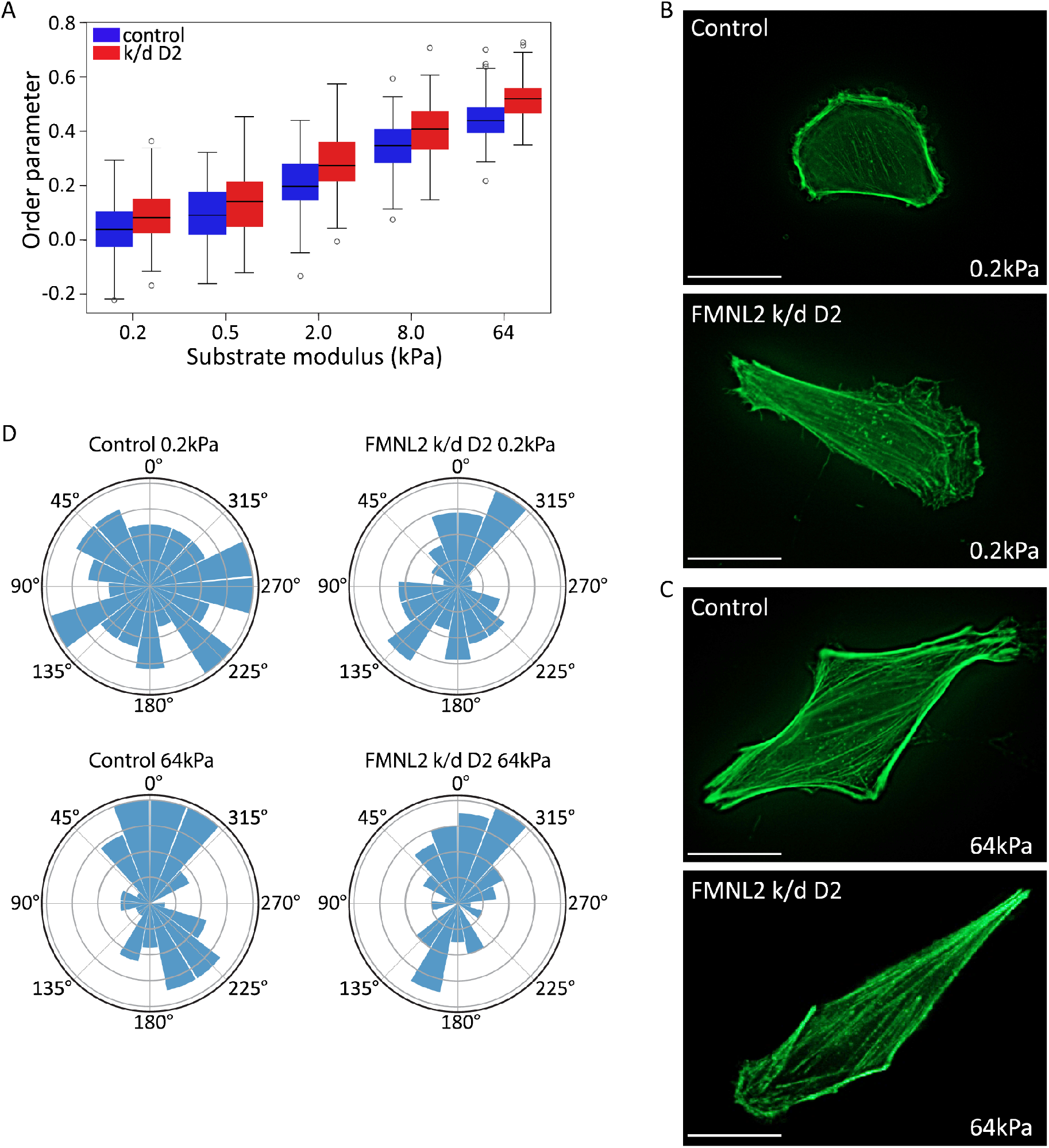
Substrate modulus affects actin stress fiber orientation in A2058 cells. (A) 2D orientational order parameter *S* quantifying average orientation of F-actin with respect to the cell Feret diameter. Increasing substrate stiffness increases the 2D order parameter. FMNL2 knockdown results are shown for FMNL2 depletion using siRNA duplex 2. Controls cells are as shown in Figure 2. ° indicates values outside of 2 standard deviations. N=3. (B, C) Representative images of the cells analyzed in (A) for control and FMNL2 depleted cells plated on the indicated substrates. Cells were fixed and stained with phalloidin to visualize actin filaments. Scale bar=20 *µ*m. (D) Windrose plots showing the orientation of the actin fibers with respect to the cells’ Feret diameter for control and FMNL2 depleted cells plated on 0.2kPa and 64kPa substrates. Increases in substrate modulus and FMNL2 depletion both yield a tighter distribution around 0, indicating increased filament alignment.

**FIG. S3:**
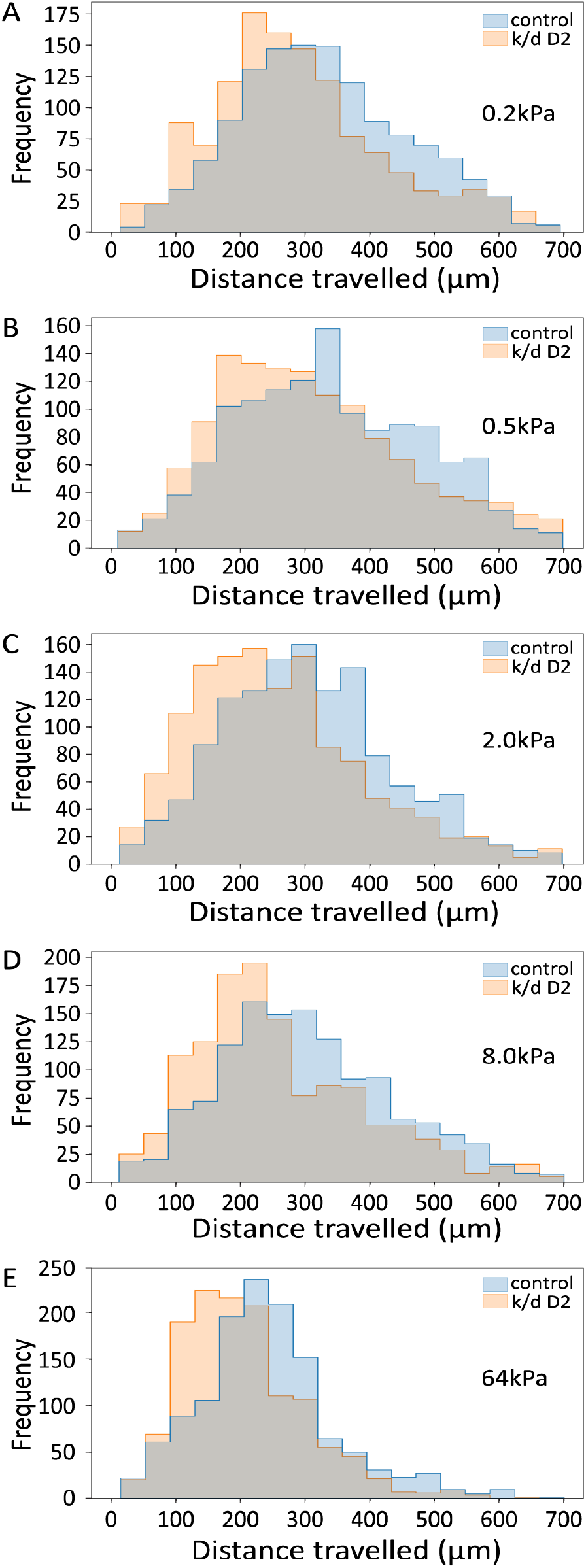
Cell motility is decreased with increased modulus. (A-E) Histograms of the net distance *D* travelled across all substrates for control (blue) and FMNL2 knockdown (siRNA duplex D2, orange) A2058 cells, with the overlap region shown in gray. Knockdown cells show a smaller average distance travelled than control cells as indicated by the shift in the histogram. Values are derived from cells tracked by live-cell imaging as with the experiments shown in Figure 1.

**FIG. S4:**
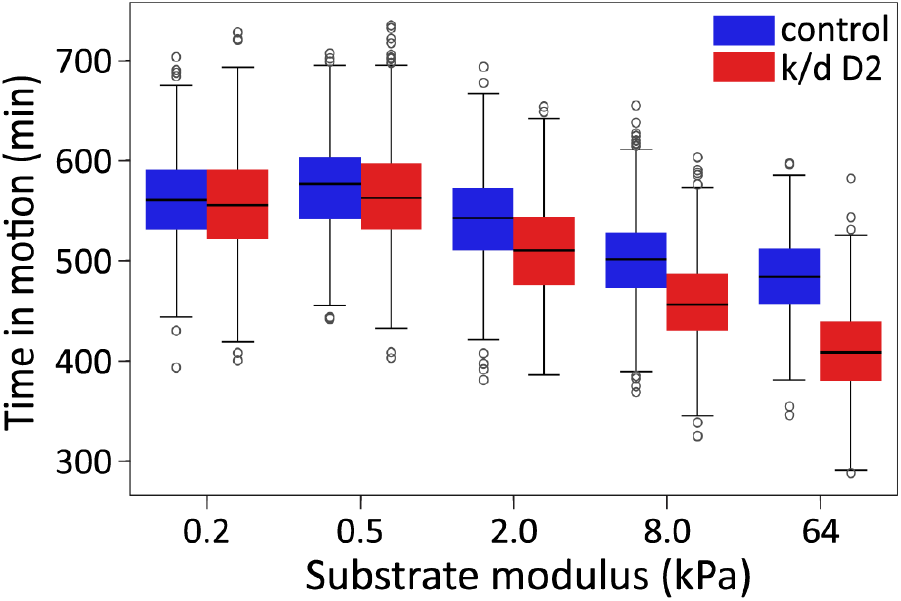
Time spent in motion for control and FMNL2 knockdown cells. (A) A box plot showing the time spent moving for control (blue) and FMNL2 knockdown (siRNA duplex D2, red) cells with increasing substrate modulus. Increasing stiffness resulted in decreased time in motion. FMNL2 depleted cells spent less time in motion than control cells on each substrate. N=3.

**TABLE S1:** Morphological changes in control and 2^nd^ duplex FMNL2 depleted A2058 cells on increasing moduli substrates. The average area (*A*), perimeter (*P*), Feret diameter (*FD*), and Roundness (*R*) are shown for control and FMNL2 knockdown cells using the 2^nd^ siRNA duplex on each substrate. The error on each value is the standard error of the mean

| Modulus<br>(kPa) | Control |  |  |  | k/d D2 |  |  |  |
| --- | --- | --- | --- | --- | --- | --- | --- | --- |
| | $A$ ($\mu\text{m}^2$) | $P$ ($\mu\text{m}$) | $FD$ ($\mu\text{m}$) | $R$ | $A$ ($\mu\text{m}^2$) | $P$ ($\mu\text{m}$) | $FD$ ($\mu\text{m}$) | $R$ |
| 0.2 | 951 ± 8 | 144.9 ± 0.8 | 44.1 ± 0.2 | 0.763 ± 0.002 | 915 ± 8 | 148 ± 1 | 46.2 ± 0.1 | 0.729 ± 0.002 |
| 0.5 | 947 ± 9 | 143.7 ± 0.7 | 44.2 ± 0.2 | 0.742 ± 0.002 | 920 ± 10 | 149.5 ± 0.8 | 47.2 ± 0.1 | 0.697 ± 0.003 |
| 2.0 | 1030 ± 8 | 154.7 ± 0.7 | 47.5 ± 0.1 | 0.704 ± 0.001 | 958 ± 8 | 163 ± 1 | 52.6 ± 0.2 | 0.638 ± 0.003 |
| 8.0 | 1050 ± 8 | 165 ± 0.8 | 52.9 ± 0.1 | 0.663 ± 0.001 | 987 ± 9 | 171.1 ± 0.9 | 56.8 ± 0.2 | 0.591 ± 0.002 |
| 64.0 | 1080 ± 7 | 175 ± 0.8 | 56.5 ± 0.2 | 0.615 ± 0.001 | 1017 ± 6 | 183 ± 0.7 | 59.6 ± 0.2 | 0.541 ± 0.001 |

**TABLE S2:**
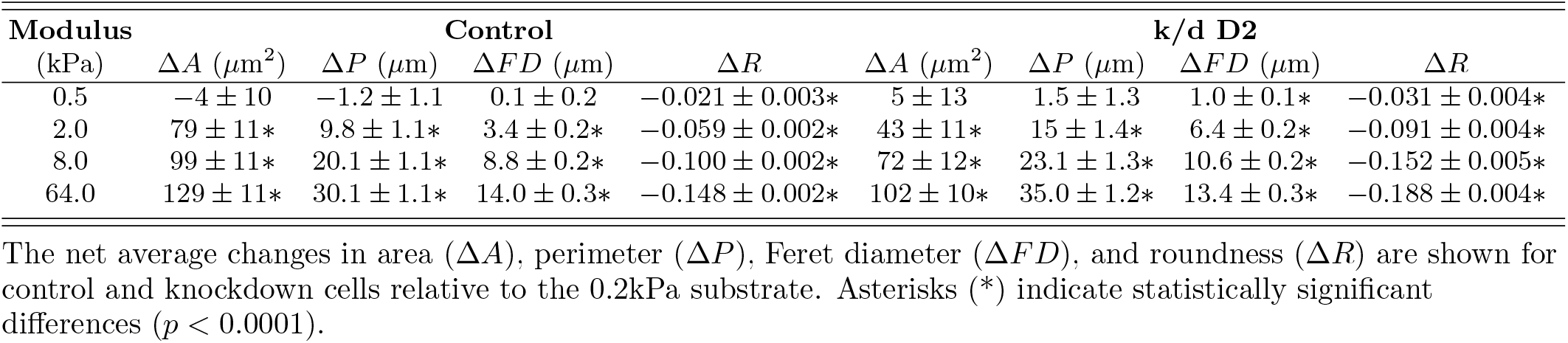
Net morphological changes between control and 2^nd^ duplex A2058 cells relative to the 0.2kPa substrate, \**p <* 0.0001.

**TABLE S3:**
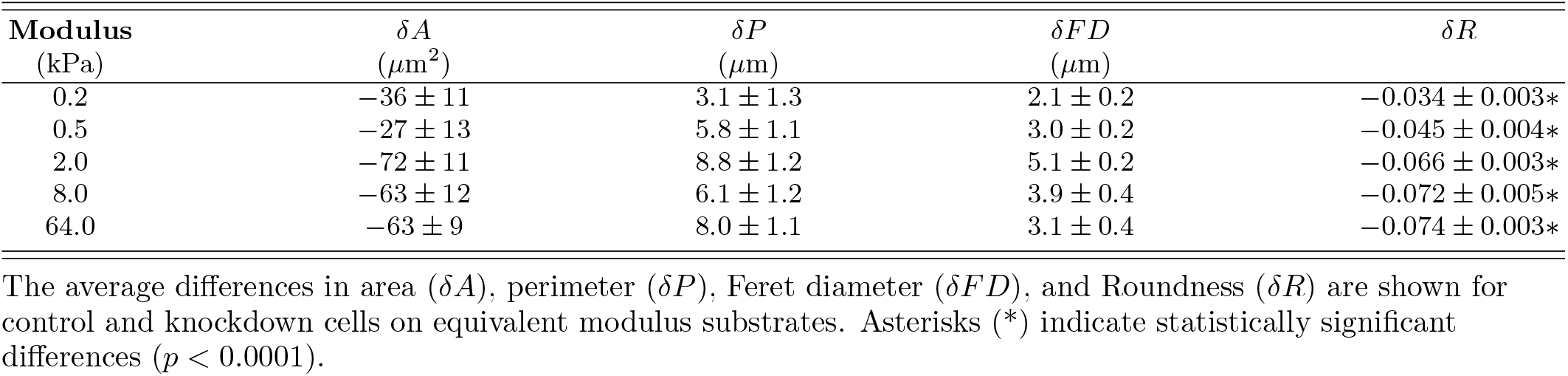
Net morphological changes between A2058 control and 2^nd^ duplex FMNL2 depleted cells on equivalent moduli substrates, \**p <* 0.0001.

**TABLE S4:** The average net distance travelled for control and 2^nd^ duplex FMNL2 depleted cells, \**p <* 0.0001.

| **Modulus**<br>(kPa) | **Control**<br>$\bar{D}$ ($\mu\text{m}$) | **k/d D2**<br>$\bar{D}$ ($\mu\text{m}$) | **Difference**<br>$\Delta\bar{D}$ ($\mu\text{m}$) |
| --- | --- | --- | --- |
| 0.2 | $329 \pm 4$ | $290 \pm 4$ | $-39 \pm 6^*$ |
| 0.5 | $340 \pm 4$ | $312 \pm 4$ | $-28 \pm 6^*$ |
| 2.0 | $303 \pm 4$ | $257 \pm 4$ | $-46 \pm 6^*$ |
| 8.0 | $282 \pm 3$ | $261 \pm 4$ | $-21 \pm 5^*$ |
| 64.0 | $242 \pm 3$ | $205 \pm 3$ | $-37 \pm 4^*$ |

Average net distance travelled ( $\bar{D}$ ) for control and knockdown cells and their relative values ( $\Delta\bar{D}$ ) on equivalent substrates. Asterisks (\*) indicate statistically significant differences ( $p < 0.0001$ ).

**TABLE S5:** The average time spent in motion for control and 2^nd^ duplex FMNL2 depleted cells, \*\*\**p <* 0.001, \*\*\*\**p <* 0.0001.

| **Modulus**<br>(kPa) | **Control**<br>$t_m$ (min) | **k/d D2**<br>$t_m$ (min) | **Difference**<br>$\Delta t_m$ (min) |
| --- | --- | --- | --- |
| 0.2 | $561 \pm 2$ | $556 \pm 2$ | $-5 \pm 3$ |
| 0.5 | $574 \pm 2$ | $564 \pm 2$ | $-10 \pm 3^{***}$ |
| 2.0 | $543 \pm 2$ | $510 \pm 2$ | $-33 \pm 3^{****}$ |
| 8.0 | $501 \pm 2$ | $458 \pm 2$ | $-43 \pm 2^{****}$ |
| 64.0 | $483 \pm 2$ | $410 \pm 1$ | $-73 \pm 2^{****}$ |

Values for the average time spent moving ( $t_m$ ) for control and knockdown cells and the difference in the average time spent moving ( $\Delta t_m$ ) between control and knockdown cells on the same substrate. \*\*\* $p < 0.001$ , \*\*\*\* $p < 0.0001$ .

**TABLE S6:** The average moving speed for control and 2^nd^ duplex FMNL2 knockdown cells, \*\*\*\**p <* 0.0001.

| **Modulus**<br>(kPa) | **Control**<br>$s_m$ ($\mu\text{m}/\text{h}$) | **k/d D2**<br>$s_m$ ($\mu\text{m}/\text{h}$) | **Difference**<br>$\Delta s_m$ ($\mu\text{m}/\text{h}$) |
| --- | --- | --- | --- |
| 0.2 | $35.5 \pm 0.2$ | $34.6 \pm 0.3$ | $-0.9 \pm 0.4^{****}$ |
| 0.5 | $36.1 \pm 0.3$ | $35.2 \pm 0.3$ | $-0.9 \pm 0.4^{****}$ |
| 2.0 | $32.7 \pm 0.3$ | $31.2 \pm 0.3$ | $-1.5 \pm 0.4^{****}$ |
| 8.0 | $33.6 \pm 0.3$ | $29.7 \pm 0.3$ | $-3.9 \pm 0.4^{****}$ |
| 64.0 | $29.6 \pm 0.2$ | $27.1 \pm 0.3$ | $-2.5 \pm 0.4^{****}$ |

Average cell speed ( $s_m$ ) for control and knockdown cells and the difference in the average cell speed ( $\Delta s_m$ ) between control and knockdown cells on the same substrate. \*\*\*\* $p < 0.0001$ .

